# Ordinary Differential Equations in Cancer Biology

**DOI:** 10.1101/071134

**Authors:** Margaret P. Chapman, Claire J. Tomlin

## Abstract

Ordinary differential equations (ODEs) provide a classical framework to model the dynamics of biological systems, given temporal experimental data. Qualitative analysis of the ODE model can lead to further biological insight and deeper understanding compared to traditional experiments alone. Simulation of the model under various perturbations can generate novel hypotheses and motivate the design of new experiments. This short paper will provide an overview of the ODE modeling framework, and present examples of how ODEs can be used to address problems in cancer biology.

## I. Introduction

Mathematical modeling of biological systems is a powerful tool to examine and investigate natural phenomena more deeply compared to using traditional experimental methods alone. Models can facilitate comprehensive qualitative analysis of biological systems, predict behavior in response to various perturbations, and motivate the design of new experiments to test the predictions. Modeling naturally belongs to the process of scientific discovery, as it leads to further biological insight while guiding the direction of future research. Many modeling frameworks have been used to address a variety of biological questions. This short paper will focus on the use of ordinary differential equations (ODEs) as tools to study the dynamics of biological systems, with specific application to cancer biology.

## II. Mathematical Overview

Ordinary differential equations describe how properties of a real-world system evolve over time. The properties are called the *state* of the system, and are chosen depending on the application at hand. For example, if we wish to understand how protein expression levels evolve in a cell line, it would be reasonable to define the state as a vector of protein expression levels. An ODE has the following general form,

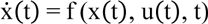

where x(t) ∈ ℝ^n^ is the *state* of the system at time t, n is the number of states, and 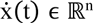 is the time-derivative of x(t), 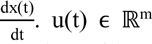 is the input vector at time t, and m is the number of inputs. The function f contains the mathematical rules which govern how x changes over time. For simplicity, we usually express x(t) as x (and u(t) as u) since dependence on time is implied. Continuing our example of protein expression dynamics, x(t) = [x_1_(t) x_2_(t) … x_n_(t)]^T^, where each x_i_(t) represents how a protein concentration of interest varies over time. u(t) may correspond to mutations in a cell line or treatments applied to a cell line. f(x(t), u(t), t) governs how the protein concentrations of interest evolve in response to the mutations or treatments. In many applications, the input u is not present or not modeled, and we simply study evolution of 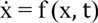.

## III. Quick Guide for Using Odes

An ODE framework is useful when the general structure of f is known from physics or other scientific principles. For instance, biochemical reactions can be modeled using mass action kinetics or Hill functions [1, 7]. Once the structure of f is chosen, experimental observations can be used to estimate the values of unknown parameters in f. Estimation of model parameters for nonlinear f is an open research challenge: collecting measurements at many time points is laborious, noise can confound true observations, and analytical expressions for parameters associated with nonlinear dynamics rarely exist. Conversely, if f is linear and a sufficient number of experiments are conducted, analytical solutions can be found using classical linear algebra approaches, such as least-squares optimization.

In practice, the number of experimental observations are often limited. Hence, many sets of parameters may describe the data equally well, meaning that specific parameter values are not particularly meaningful. Instead, *relative* parameter values and *qualitative* trends can improve understanding of the interactions and mechanisms that govern biological system behavior. In addition, *sensitivity* analysis should be conducted in order to determine how parameter values vary with small perturbations in the experimental data; useful models tend to be less sensitive, i.e., robust, to small changes in data.

Once a set of model parameters are determined from experimental data (i.e., f is fully specified), one should simulate the model and compare simulated behavior with experimental data. Often, one reserves a portion of data for model fitting (i.e., determining f) and the remaining data for model testing (i.e., comparing simulated and experimental results). This procedure is known as *cross-validation*, and provides a measure of modeling accuracy and limitations. A suitable model will generally attain qualitative agreement between simulations and data.

Moreover, the model parameters should be analyzed from a qualitative perspective to improve understanding of the biological system dynamics. For example, if the parameters were reaction rates, parameter ratios would describe relative frequencies of the reactions.

Further, the ODE can be simulated under new conditions to generate novel hypotheses that drive future experiments. The following section will show how an ODE model was used to propose a new chemotherapy treatment strategy, driven by promising results in simulation.

## IV. Application to Cancer Biology: Case Study

Below we describe the work of Itani et al. as a noteworthy example of using an ODE model for a cancer biology application. In this research, an ODE model was utilized to understand the effects of chemotherapy drugs applied to breast cancer cells and to propose a new time-dependent treatment strategy [1].

### A. Background

Itani et al. proposed an ODE model of a signaling pathway in breast cancer cell lines that overexpress HER2 [1]. HER stands for human epidermal growth factor receptor, and is known to play a significant role in the progression of cancer [2]. The model was based on experimental observations from a leading *Nature* study, Sergina et al., which revealed how HER2-positive breast cancer cells escape inhibition by promising anticancer drugs, called tyrosine kinase inhibitors (TKIs) [1, 3]. Sergina et al. discovered that a shift in the HER3 phosphorylation-dephosphorylation equilibrium, driven by Akt negative feedback signaling, causes TKIs to be ineffective in HER2-driven breast cancers [3].

### B. Construction and use of the ODE model

The modeling goal in Itani et al. was to choose a sufficiently simple representation of the signaling pathway involving HER2, HER3 and Akt that reproduced the experimental observations of [3] and facilitated the design of novel treatment strategies [1]. Signal transmission via phosphorylation of proteins was represented using ODEs, assuming first- and second-order mass action kinetics [1]. For example, the equation associated with protein A has the form,

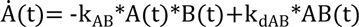

where k_AB_ is the rate of binding of protein A to protein B and k_dAB_ is the rate of auto-decomposition of protein complex AB [1]. Moreover, trafficking of HER3 between the cytoplasm and cell membrane was modeled using an ODE framework that abstracted these two regions as compartments, and utilized special variables to transfer HER3 between the compartments [1].

A (non-unique) parameter set that produced qualitative agreement between model simulations and experimental observations of [3] was determined using a search algorithm [1]. The model was then used to explore the effects of various treatment strategies and identify one that showed improved performance over the standard approach, simple application of a TKI, in simulation [1]. Experimental validation *in vitro/vivo* is still needed and reserved for future work [1].

### C. Discussion

The proposed treatment scheme of [1] was *sequential*, meaning that certain drugs were applied in a specially-timed sequence to produce a desired outcome. Understanding the signaling pathway *dynamics* (i.e., how protein concentrations change over time) was essential for determining this time-dependent strategy. Construction of the ODE dynamical system model facilitated the investigation of new treatment strategies in simulation, without the use of laborious experiments until a promising approach was identified. Thus, the model allowed for *intelligent experiment design*, whereas biological research without mathematical models must rely on intuition and guesswork to guide the development of future experiments.

## V. Additional Illustrative Examples

In the following section, we briefly present three additional examples of using ODEs in cancer biology research. In each work, an ODE model was estimated from experimental observations to represent a system of cancer cells, and supported existing knowledge or contributed to new biological insight.

Gupta et al. studied how phenotypes of cancer cells transition naturally and under therapeutic stress using a special type of linear ODE, called a *Markov model* [4]. The model represented changes in cellular phenotype as *stochastic* transitions in discrete time [4]. Interestingly, the model predicted that cancer non-stem-like cells can transition into stem-like cells (at a low rate), which counters the classical unidirectional view of stem-like cells [4]. The major contribution of this work is a systematic procedure for characterizing how treatments influence phenotype transitions in cancer cell populations, using data collected at only two time points [4].

Moreover, Goldman et al. also used an ODE model to represent phenotype transitions in response to anticancer drugs [5]. A linear time-invariant ODE was constructed to theoretically test drug-induced phenotypic plasticity versus clonal selection [5]. The model described how the quantity of cells in each phenotype changed over time using parameters, such as net reproductive rates and phenotype switching rates [5]. The mathematical model predicted that chemotherapy-tolerant cells arise from non-cancer-stem-like cells, which was later verified in experiments [5]. This contributed to the main conclusion of the paper that cancer treatment induces phenotype transition into a chemotherapy-tolerant state [5].

In a final example, Dobbe et al. developed a methodology to estimate a linear time-varying ODE model of the signaling network in HER2 breast cancer cell lines [6]. Measurements of protein expression levels (from several cell lines with various treatments applied) were formulated into a convex optimization problem [6]. An ODE model describing the protein expression dynamics was obtained from the solution [6]. In addition, existing biological knowledge was identified in the solution [6]. This work provides a systematic mathematical framework for studying heterogeneity in cancer, as analysis of data sets composed of many treatments and cell types is often infeasible without computational tools.

## VI. Conclusions

Ordinary differential equation-based models are useful in cancer biology to study how biological systems change over time. Collecting data at *several* time points is essential for estimating sufficiently accurate models. Models should be analyzed from a qualitative point of view to obtain deeper understanding of the underlying dynamics. Here we presented several examples of using ODEs in cancer biology applications. ODEs, and mathematical models in general, can support experimental findings and lead to new avenues of scientific discovery.

## References

[1] Itani, S., Gray, J., & Tomlin, C. J. (2010, June). An ODE Model for the HER2/3-AKT Signaling Pathway in Cancers that Overexpress HER2. In American Control Conference (ACC), 2010 (pp. 1235–1241). IEEE.

[2] Slamon, D. J., Clark, G. M., Wong, S. G., Levin, W. J., Ullrich, A., & McGuire, W. L. (1987). Human breast cancer: correlation of relapse and survival with amplification of the HER-2/neu oncogene. Science, 235(4785), 177–182.

[3] Sergina, N. V., Rausch, M., Wang, D., Blair, J., Hann, B., Shokat, K. M., & Moasser, M. M. (2007). Escape from HER-family tyrosine kinase inhibitor therapy by the kinase-inactive HER3. Nature, 445(7126), 437–441.

[4] Gupta, P. B., Fillmore, C. M., Jiang, G., Shapira, S. D., Tao, K., Kuperwasser, C., & Lander, E. S. (2011). Stochastic State Transitions Give Rise to Phenotypic Equilibrium in Populations of Cancer Cells. Cell, 146(4), 633–644.

[5] Goldman, A., Majumder, B., Dhawan, A., Ravi, S., Goldman, D., Kohandel, M., Majumder, P. K., & Sengupta, S. (2015). Temporally sequenced anticancer drugs overcome adaptive resistance by targeting a vulnerable chemotherapy-induced phenotypic transition. Nature communications, 6.

[6] Dobbe, R., Chang, Y. H., Korkola, J., Gray, J., & Tomlin, C. J. (2015, July). Heterogeneity in cancer dynamics: A convex formulation to dissect dynamic trajectories and infer LTV models of networked systems. In American Control Conference (ACC), 2015 (in press). IEEE.

[7] Chang, Y. H., Dobbe, R., Bhushan, P., Gray, J. W., & Tomlin, C. J. (2014). Reconstruction of Gene Regulatory Networks based on Repairing Sparse Low-rank Matrices. bioRxiv, 012534.

